# Combined PI3K and MAPK inhibition synergizes to suppress PDAC

**DOI:** 10.1101/2023.08.15.553438

**Authors:** Bailey A. Bye, Jarrid Jack, Alexandra Pierce, R. McKinnon Walsh, Austin Eades, Prabhakar Chalise, Appolinaire Olou, Michael N. VanSaun

## Abstract

Oncogenic KRAS mutations are nearly ubiquitous in pancreatic ductal adenocarcinoma (PDAC), yet therapeutic attempts to target KRAS as well as its target MAPK pathway effectors have shown limited success due to the difficulty to pharmacologically target KRAS, inherent drug resistance in PDAC cells, and acquired resistance through activation of alternative mitogenic pathways such JAK-STAT and PI3K-AKT. While KRAS canonically drives the MAPK signaling pathway via RAF-MEK-ERK, it is also known to play a role in PI3K-AKT signaling. Our therapeutic study targeted the PI3K-AKT pathway with the drug Omipalisib (p110α/β/δ/γ and mTORC1/2 inhibitor) in combination with MAPK pathway targeting drug Trametinib (MEK1/2 inhibitor) or SHP099-HCL (SHP099), which is an inhibitor of the KRAS effector SHP2. Western blot analysis demonstrated that application of Trametinib or SHP099 alone selectively blocked ERK phosphorylation (pERK) but failed to suppress phosphorylated AKT (pAKT) and in some instances increased pAKT levels. Conversely, Omipalisib alone successfully inhibited pAKT but failed to suppress pERK. Therefore, we hypothesized that a combination therapeutic comprised of Omipalisib with either Trametinib or SHP099 would inhibit two prominent mitogenic pathways, MEK and PI3K-AKT, to more effectively suppress pancreatic cancer. *In vitro* studies demonstrated that both Omipalisib/Trametinib and Omipalisib/SHP099 combination therapeutic strategies were generally more effective than treatment with each drug individually at reducing proliferation, colony formation, and cell migration compared to vehicle controls. Additionally, we found that while combination Omipalisib/SHP099 treatment reduced implanted tumor growth *in vivo*, the Omipalisib/Trametinib treatment was significantly more effective. Therefore, we additionally tested the Omipalisib/Trametinib combination therapeutic in the highly aggressive PKT (Ptf1a^cre^, LSL-Kras^G12D^, TGFbR2^fl/fl^) spontaneous mouse model of PDAC. We subsequently found that PKT mice treated with the Omipalisib/Trametinib combination therapeutic survived significantly longer than mice treated with either drug alone, and more than doubled the mean survival time of vehicle control mice. Altogether, our data support the importance of a dual treatment strategy targeting both MAPK and PI3K-AKT pathways.

## Introduction

Pancreatic ductal adenocarcinoma (PDAC) is one of the deadliest cancers with a 5-year survival rate of 11%, due in part to inherent and acquired resistance to limited therapeutic options (1). Gemcitabine has been the standard first-line chemotherapeutic for PDAC for more than 20 years, but only increases patient survival by 6-7 months (2, 3). Other, more aggressive treatment options, such as FOLFIRINOX, only increase median overall survival by 4-5 months compared to Gemcitabine (3, 4). Difficulty effectively suppressing PDAC tumor growth with chemotherapy is a reflection of, among other things, mutations harbored by the cancer cells themselves. KRAS is mutated in over 90% of PDAC, with the G12D mutation being the most common (5-8). In a normal cell, KRAS functions as a binary signaling switch by conversion from an inactive, guanosine diphosphate (GDP)-bound conformation to an active guanosine triphosphate (GTP)-bound conformation, which can then activate multiple downstream signaling cascades (9). Constitutive activation of KRAS as a result of G12D mutation impairs GTPase function via steric hindrance of GTP hydrolysis, which prolongs KRAS in the active GTP-bound state and leads to increased activation of downstream mitogenic pathways (10, 11). The high frequency of mutated KRAS in PDAC has driven interest for targeted therapeutic strategies. However, while a drug targeting KRAS G12C has been approved by the FDA to treat non-small cell lung cancer (12), this mutation is present in less than 3% of PDAC (13). Many of the other drugs targeting mutated KRAS or its downstream effectors including ERK, MEK, and AKT have so far proven ineffective as individual therapeutic targets (6). Alternatively, targeting upstream of mutant KRAS via effectors such Src homology phosphatase 2 (SHP2) has been the focus of recent studies (14-16).

The MAPK and PI3K-AKT pathways are a central focus for investigation in PDAC due to their prominent roles in cell proliferation and survival and the fact that they are known downstream targets of mutant KRAS (17-19). A highly potent and specific MEK1/2 (MAPK pathway) inhibitor, Trametinib, has been approved by the FDA to treat advanced melanoma tumors, harboring a BRAF V600E mutation, in combination with the BRAF inhibitor dabrafenib (20-25). MEK inhibitors such as trametinib have shown promise as therapeutic strategies for PDAC, but display a limited efficacy in clinical trials due in part to the development of resistance (26-29). To overcome resistance to MEK inhibition, many current studies are focused on combined therapeutics to block reactivation pathways (30, 31).

Another strategy for targeting mutant KRAS-driven MAPK activation is inhibiting KRAS interacting proteins, such as the tyrosine phosphatase SHP2, which is an essential factor for RAS-mediated MAPK signaling (32-34). Recently, SHP2 was shown to be required for mutant KRAS-driven PDAC tumorigenesis (34). Targeting SHP2 via a selective inhibitor such as SHP099 (35) effectively inhibits the MAPK pathway and counteracts therapeutic resistance when combined with a MEK inhibitor in mutant KRAS-driven cancers (14-16). While MAPK inhibition via SHP099 in combination with PI3K inhibitors has shown efficacy in other types of cancer (36, 37), it remains to be investigated in PDAC. The drug Omipalisib is a potent inhibitor of PI3K complex components p110α/β/δ/γ and downstream factors mTORC1/2, ultimately suppressing activation of AKT (38). Combined MEK inhibition and PI3K pathway inhibition (via AKT or mTOR) have shown effectiveness in PDAC as well as other cancers (18, 26, 39-44). While MAPK inhibition via SHP099 in combination with PI3K inhibitors has shown efficacy in a few cancers (36, 37), it remains to be investigated in PDAC. Omipalisib has been investigated clinically as a single therapeutic agent in solid tumors, but its efficacy was modest, suggesting that further investigation into an optimal combination treatment strategy is needed (45).

The purpose of our current study was to test the efficacy of targeting multiple aspects of MAPK inhibition via SHP099 or Trametinib combined with PI3K pathway inhibition via Omipalisib in PDAC. Our results identified a persistence of PI3K-AKT pathway activation upon MAPK pathway inhibition via Trametinib or SHP099. Subsequently, a greater suppression of *in vitro* tumor cell proliferation and migration *in vitro* as well as tumor size *in vivo* was observed with dual PI3K/MAPK inhibition.

## Materials and Methods

### Cell culture and reagents

Mouse PDAC tumor cell lines K8484 was derived from KPC (Ptf1a^cre/+^;LSL-Kras^G12D/+^) and PKT62 was derived from PKT (Ptf1a^cre/+^;LSL-KRAS^G12D/+^; TGFBR2^flox/flox^) genetically-engineered mouse models of PDAC. These cell lines were kindly gifted by Dr. David Tuveson (K8484) (46) or Dr. Nipun B. Merchant / Dr. Nagaraj S. Nagathihalli (PKT62). MiaPaCa-2 and Panc1 are primary human PDAC tumor cell lines purchased from ATCC. All cell lines were maintained in high-glucose (4.5 g/L) Dulbecco’s Modified Eagle Medium (Gibco, #1-995-073) supplemented with 5% heat-inactivated fetal bovine serum (Biowest, #S1260) and antibiotic-antimycotic (Gibco, #15240062) at 37°C with 5% CO_2_. Media was refreshed every 2-3 days. Omipalisib (p110α/β/γ/δ inhibitor; catalytic subunit of PI3K; cat# S2658), Trametinib (MEK 1/2 inhibitor, cat# S2673), and SHP099-HCl (SHP2 inhibitor, cat#S8278) were purchased from Selleckchem (Houston, TX, USA). For *in vitro* drug experiments, DMSO controls matched the DMSO concentration in wells containing a combination therapeutic treatment. Unless otherwise specified, drug treatments were diluted to final concentration in DMEM + 5% FBS and, in experiments lasting longer than 48 hours, treatment media was refreshed every 2 days.

### Western blot and antibodies

Cells were lysed in cold RIPA buffer (Cell Signaling #9806S) supplemented with NaF (10mM), sonicated, then centrifuged for 10 minutes at 12,000 x g. Tumor tissue collected from *in vivo* studies was homogenized in NaF supplemented RIPA buffer before sonication and centrifugation. Protein concentration in each lysate was determined using a Pierce BCA protein assay (Thermo Scientific #PI23227). Lysate samples of equal protein concentration were loaded and run on a 10% SDS-PAGE gel before transfer to nitrocellulose membranes. Membranes were blocked in TBS with 2% milk and 2% BSA before incubation with primary antibodies at 4°C overnight followed by incubation with appropriate secondary antibodies for 1 hour at room temperature. Primary antibodies against phospho-AKT (S473, #4060), total AKT (pan, #4298), phospho-p44/42 MAPK (Erk1/2, Thr202/Tyr204, #4370), p44/42 MAPK (Erk1/2, #46953), β-Actin (#CS12262) were purchased from Cell Signaling. Secondary antibody, peroxidase-conjugated anti-rabbit IgG (#711036152) was from Jackson Immunoresearch. If applicable, blots were stripped using stripping buffer (Thermo Scientific, #PI34577) for 30 minutes at room temperature and imaged to verify complete stripping of antibody. Pico or femto substrate (Thermo Scientific #PI34577 or #PI34095) was added to all blots prior to imaging using a Fluorchem M imager (Bio-Techne, 92-15312-00).

### EdU proliferation assay

Measuring cell proliferation via 5-ethynyl-2’-deoxyuridine (EdU) incorporation has been previously described (47). Briefly, cells were seeded at 50k cells/well in a 24 well plate and allowed to attach overnight. The next morning, drug treatments were applied to respective wells. After 24 hours, wells were spiked with EdU (Invitrogen, A10044) to a final concentration of 20µM and allowed to incubate for 6 hours. Cells were then detached from the plate with trypsin (Gibco, #25-200-056) and fixed overnight on a rocker at 4°C in 5% buffered formalin. After fixing, cells were permeabilized and incubated with incubated with copper sulfate, azide dye (AF647, Invitrogen, #AI0277), sodium ascorbate and stained with PI (0.5µg/mL) (Invitrogen, #P3566) before analysis via flow cytometry. Cells were gated for singlets, then PI positivity, then EdU positivity. Cell proliferation was reported as percentage of EdU+ cells within the total PI positive cells, using negative control cells (not incubated with EdU) to define the gating for analysis.

### Migration assay

Cells were first seeded in a confluent layer on a 24-well plate (300-400k cells/well, depending on the cell line) and allowed to attach overnight. The next morning, the cells were incubated with Mitomycin C (5µg/mL) (Thermo Scientific, #AAJ67460XF) for 2 hours to halt proliferation (excluding the PKT cell line, since the migration assay was conducted over only 8 hours). A scratch was then created across the center of each well with a sterile pipette tip and each well was washed with PBS to remove debris. Treatment media (drug in DMEM + 2.5% FBS) was then added and each well was imaged to provide a baseline for migration calculations (T0). Endpoint images were collected at 8 (PKT62), 24 (Panc1 and K84848 OmiTram), and 48 (MiaPaCa and K8484 OmiSHP) hours after scratch was created. Percent wound closure was calculated using ImageJ.

### Colony formation assay

Cells were seeded at 1000 cells/well in a 6 well plate and allowed to attach overnight. Treatment media was then added to each well and was refreshed every two days. At endpoint, cells were stained with crystal violet solution (Sigma Aldrich, #V5265) for 20 minutes and rinsed with water. After imaging the plates, the crystal violet was dissolved in 10% acetic acid and absorbance for each well was read at 590 nm.

### Histology

Tissues were fixed immediately after collection in 10% formalin at room temperature overnight and transferred to 70% ethanol before paraffin processing, embedding, and sectioning. Slides were rehydrated and stained with either Hematoxylin and Eosin (Electron Microscopy Sciences, #26252-01 and #26252-02) or processed for immunofluorescence (IF) staining. For IF staining, tissue sections were permeabilized using TBS with 0.05% Triton X-100 (Sigma Aldrich, #T9284) for 15 minutes. Antigen retrieval was performed by microwaving slides in 10mM sodium citrate buffer (pH 6.0). Tissue sections were blocked in donkey serum-containing IF blocking buffer for 1 hour at room temperature before adding primary antibodies in a 1:1 ratio of donkey serum-containing blocking buffer and TBS for overnight incubation at 4 degrees C. Slides were then incubated with secondary antibody in blocking buffer containing (5µg/mL) Hoechst nuclear stain (Invitrogen #H3570) for 1 hour at room temperature. Slides were mounted using Invitrogen ProLong Diamond Antifade Mountant (Thermo Fisher #P36961) for IF slides and Permount mounting medium (Thermo Fisher #SP15-100) for H&E slides. All slides were imaged using an EVOS E5000 (Thermo Fisher #AMF5000) microscope. Primary antibodies used for IF Staining were Claudin 18 (Proteintech, #66167-1) and phospho-histone H3 (Cell Signaling, #9701), and secondary antibodies were Alexa-Fluor 488 donkey anti-mouse IgG (H+L) (#A21202) and Alexa-Fluor 594 donkey anti-rabbit IgG (H+L) (#A21207). For tumor area quantification, slides were scanned using a Pathscan Enabler IV slide scanner (Meyer Instruments) and normal pancreas, tumor, necrotic, and other/unspecified areas were quantified using Adobe Photoshop. Tissues from mice that unexpectedly succumbed to disease, resulting in non-viable tissue, were excluded from histological analysis.

### Mice

In accordance with approved IACUC guidelines, subcutaneous tumors were established by injecting 1 x 10^6^ tumor cells tumor cells into the flank of C57BL/6 mice. Tumor size was monitored every 2 days using calipers. In both sets of *in vivo* experiments, the subcutaneous model or PKT GEM model of PDAC, respective drugs were dissolved in 0.5% methylcellulose with >0.05% Tween80 and administered via oral gavage 3 times per week (1mg/kg Trametinib, 0.3mg/kg Omipalisib, 50 mg/kg SHP099). Treatment for mice with subcutaneous tumors began when tumors reached an approximate volume of 100 mm^3^ and continued for either 18 weeks or until tumors reached endpoint size of 2000 mm^3^. In the PKT mice, drug treatment began at 4.5 weeks of age and continued for 10 weeks or until the humane endpoint was reached. When possible, mice were euthanized 6 hours after the final dose of drug in order to verify respective targeted pathway inhibition.

## Statistical analysis

For the proliferation and migration assays, 4 replicate wells were analyzed per experimental treatment in all 4 cell lines. For the colony-forming assays, 3 replicate wells were analyzed per experimental treatment in all 4 cell lines. For the subcutaneous tumor experiments, vehicle: n=5; Omipalisib: n=6; Trametinib: n=6; SHP099: n=6; OmiTram: n=5; OmiSHP: n=6. Tumor growth among the six experimental groups of mice were measured at baseline and every two days until 18 days (0, 2, 4, 6, 8, 10, 12, 14, 16, 18). Differences in tumor growth over time among the groups were analyzed using linear mixed models for repeated measure data using SAS procedure GLIMMIX (SAS Institute Inc.) using AR(1) covariance structure. Tumor growth between groups over time interactions were assessed by specifying appropriate contrast statements within the modelling framework. For the *in vivo* experiments using PKT mice, Vehicle: n=6; Omipalisib: n=6; Trametinib: n=6; OmiTram: n=7. Survival analysis included all listed mice, but all other post-mortem analyses excluded mice that unexpectedly succumbed to disease before final weights and tissues could be collected (post-mortem analysis: vehicle: n=5; Omipalisib: n=6; Trametinib: n=5; OmiTram n=6). In all analyses except that of tumor growth rate in the *in vivo* subcutaneous tumor experiments, statistical significance was assessed by one-way ANOVA with Tukey’s multiple comparison analysis using GraphPad Prism5 software (*p < 0.05, **p <0.01, ***p < 0.001, and ****p < 0.0001).

## Results

### Inhibition of MAPK or PI3K signaling promotes alternative mitogenic signaling pathway activation

Tumor cell persistence in response to therapeutic drugs is due in part to their ability to activate alternative tumorigenic signaling pathways when other mitogenic pathways are suppressed (48). The MAPK and PI3K-AKT pathways are two prominent mitogenic pathways driving pancreatic ductal adenocarcinoma (PDAC), in part due to constitutive activation of KRAS, which is mutated in almost all PDAC (5-8). Therefore, we aimed to assess the pathway activation status of each protein after treatment with targeted therapeutics. We measured PDAC cell response to drugs targeting the MEK or PI3K-mTOR pathways using Omipalisib (PI3K pathway inhibitor), Trametinib (MAPK pathway inhibitor via ERK), and SHP099 (MAPK pathway inhibitor via SHP2). Effective drug concentrations were determined by phospho-specific detection of target proteins via dose response curves (SF1-3).

First we assessed pathway action after MEK inhibition via Trametinib in mouse (K8484, PKT62) and human (MiaPaCa2, Panc1) PDAC cell lines, which demonstrated effective ERK inhibition at 5nM (PKT62), 10 nM (K8484, MiaPaCa2), and 50nM (Panc1) (SF2 and Figure 1). Additionally, we noted sustained or upregulated AKT activation, as measured by phosphorylated AKT (pAKT) (Figure 1 and SF2). Next, we assessed ERK activation in response to SHP2 inhibition (SHP099) across all PDAC cell lines. While inhibiting pERK in PKT62, MiaPaCa2, and Panc1 cell lines, SHP099 did not appear to induce a compensatory increase in PI3K pathway activation, except in the Panc1 cell line, which showed a slight increase in pAKT at 20µM (SF3). Next, we tested whether treatment with PI3K pathway inhibitor Omipalisib would induce a similar compensatory activation of the MAPK pathway. Omipalisib demonstrated effective inhibition of AKT phosphorylation at 5nm (K8484) and 25nM (PKT62, MiaPaCa2, Panc1), and we observed sustained or upregulated MAPK activation as measured by pERK levels (Figure 1 and SF1). Together, these results indicate that PDAC cells may respond to AKT inhibition by activating ERK and vice-versa, thereby suggesting that each mitogenic pathway contributes resistance to drugs targeting the other. Therefore, we hypothesized that simultaneous MAPK/PI3K pathway inhibition via combined Trametinib and Omipalisib treatment (OmiTram) or SHP099 and Omipalisib (OmiSHP) would more effectively inhibit PDAC tumor cells than Trametinib, Omipalisib, or SHP099 alone.

**Figure 1.**
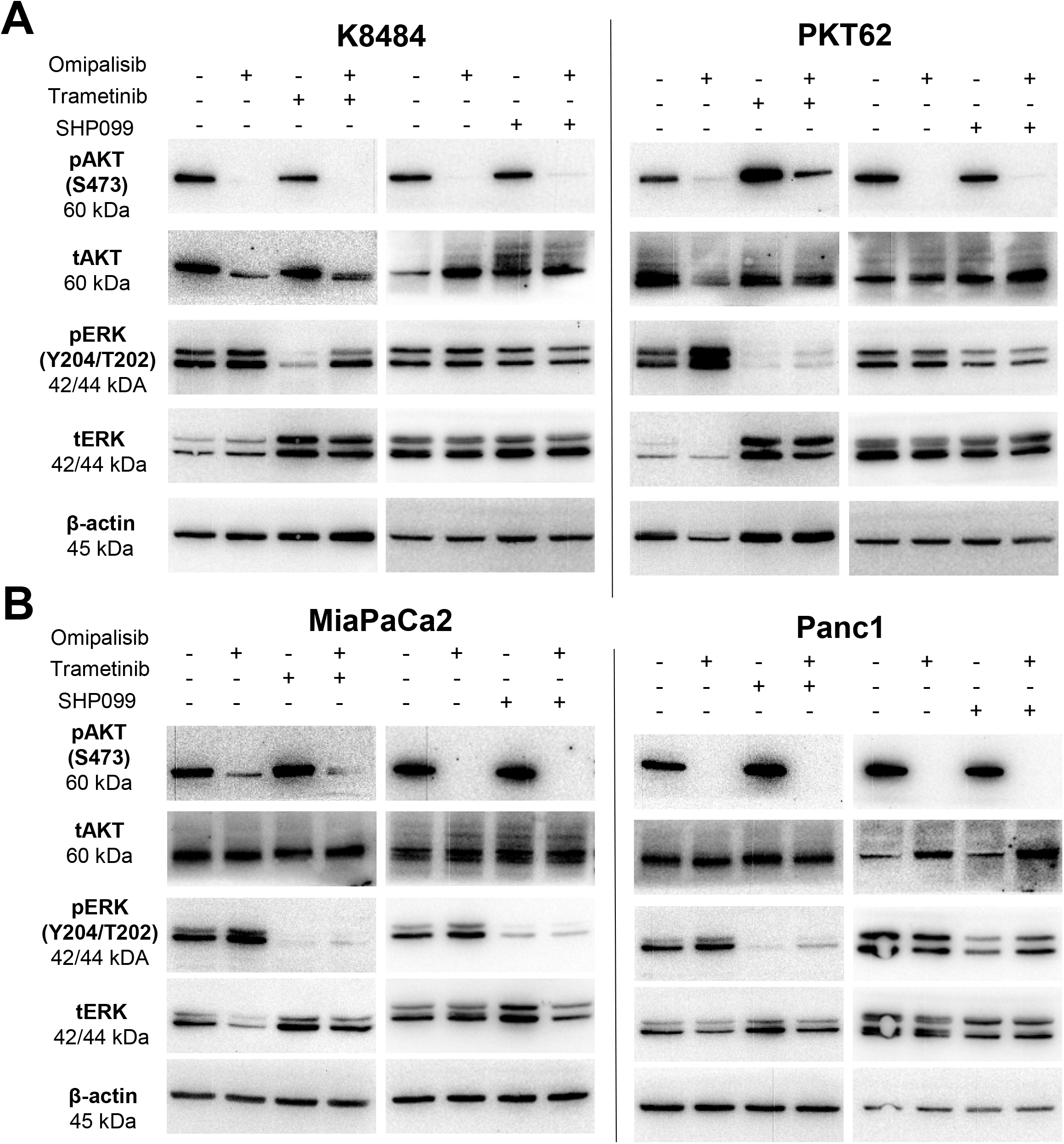
Inhibition of MAPK or PI3K signaling promotes alternative mitogenic signaling pathway activation. Western blots demonstrating inhibition and/or activation of either PI3K pathway (pAKT compared to tAKT levels) or MAPK (pERK compared to tERK levels) upon targeted therapeutic treatment for 24 hours for Omipalisib and Trametinib, and 3 hours for SHP099. **(A)** K8484 cells were treated with 5nM Omipalisib, 10nM Trametinib, and/or 50uM of SHP099. **(B)** PKT62 cells were treated with 25nM Omipalisib, 5nM Trametinib, and/or 20 uM SHP099. **(C)** MiaPaCa2 **(C)** and Panc1 **(D)** cells were treated with 25nM Omipalisib, 20 nM Trametinib, and/or 20 uM SHP099.

### Combined targeting of PI3K and MAPK pathways inhibits proliferation, colony forming ability, and migration of PDAC cells *in vitro*

Combination treatment using OmiTram suppressed pERK and pAKT in all tested cell lines (Figure 1). OmiSHP combination treatment suppressed pAKT in all cell lines and pERK in 3 of 4 cell lines (PKT62, MiaPaCa2, and Panc1), though to a lesser degree (Figure 1). To assess the effect of MAPK/PI3K inhibition on PDAC cell aggressiveness *in vitro*, we assessed the effect of OmiTram or OmiSHP treatment on PDAC tumor cell proliferation, migration, and colony-forming ability. First, we assessed OmiSHP or OmiTram effect on cell proliferation via EdU incorporation assay. Combination treatment with OmiTram more effectively suppressed cell proliferation than either Omipalisib or Trametinib treatment alone in all cell lines tested (Figure 2a). Combination treatment with OmiSHP also suppressed proliferation in all cell lines, but it was more effective than either Omipalisib or SHP099 treatment alone in two of the four cell lines tested (K8484 and Panc1) (Figure 2b).

**Figure 2.**
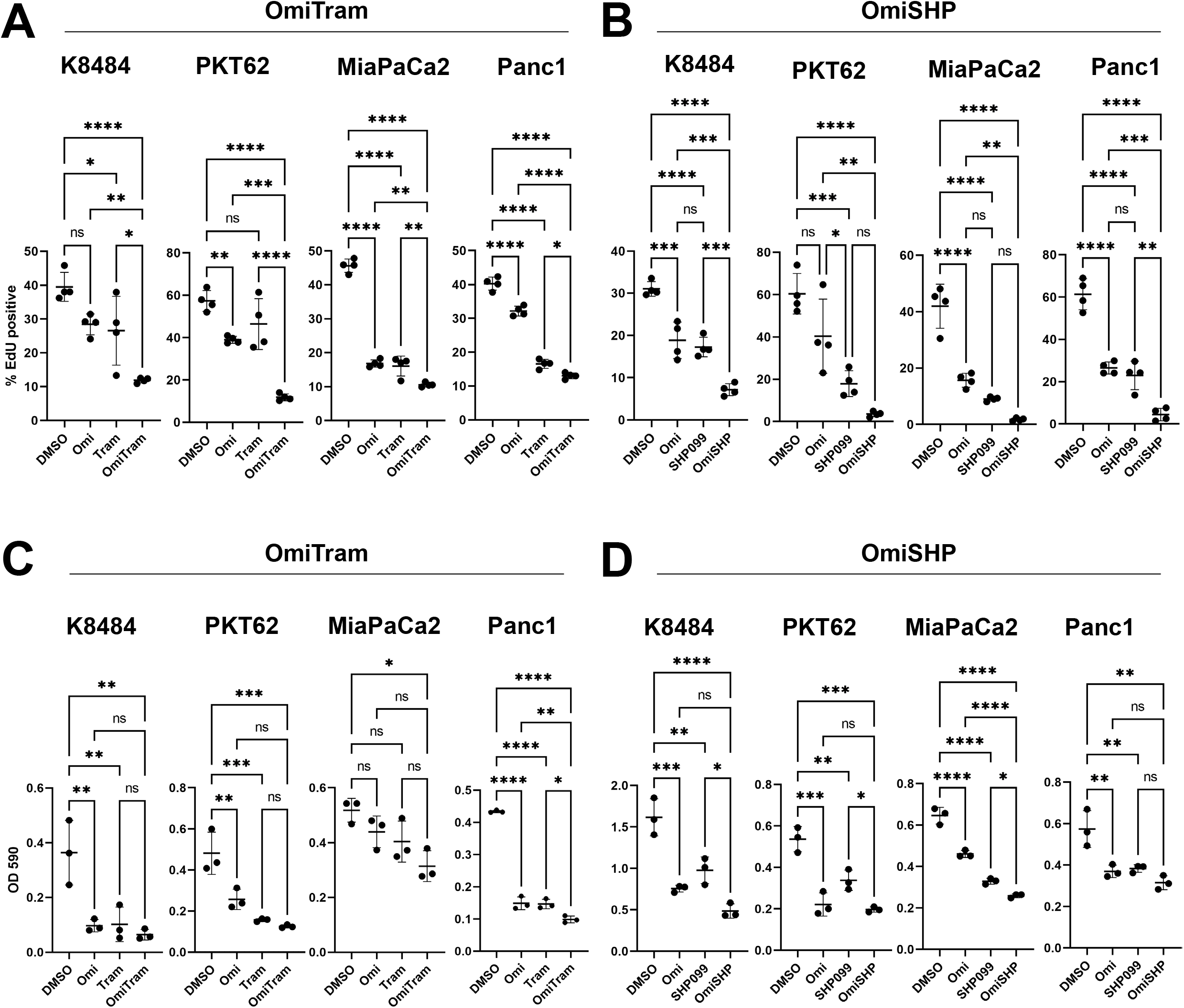
Combined targeting of PI3K and MAPK pathways inhibits proliferation and colony forming ability of PDAC cells *in vitro*. Changes in cell proliferation in response to either OmiTram **(A)** or OmiSHP **(B)** treatment for 24 hours was measured via EdU assay. K8484 cells were treated with 5nM Omipalisib, 10nM Trametinib, and/or 50 uM SHP099. PKT62 cells were treated with 25nM Omipalisib, 10 nM Trametinib and/or 20 uM SHP099. MiaPaCa2 cells were treated with 25nM Omi, 1nM Tram, and/or 20uM SHP099. Panc1 cells were treated with 25nM Omipalisib, 1nM Trametinib, and 20uM SHP099. Changes in colony-forming ability in response to either OmiTram. n=4 wells for all groups. (*p < 0.05, **p <0.01, ***p < 0.001, and ****p < 0.0001). **(C)** or OmiSHP **(D)** treatment was measured via absorbance from crystal violet stain dissolution. K8484 cells were treated with 5nM Omipalisib, 10nM Trametinib, and/or 50 uM SHP099. PKT62 cells were treated with 5nM Omipalisib, 10nM Trametinib, and/or 2 uM SHP099. MiaPaCa2 cells were treated with 5nM Omipalisib, 10nM Trametinib, and/or 20 uM SHP099. Panc1 cells were treated with 50nM Omipalisib, 20nM Trametinib, and/or 50uM SHP099.. n=3 wells for all groups. (*p < 0.05, **p <0.01, ***p < 0.001, and ****p < 0.0001).

We further tested the effect of each combination therapeutic on tumor cell survival and proliferation via colony forming assay (Figure 2c, d). The OmiTram dual therapeutic more effectively suppressed colony formation in MiaPaCa2 and Panc1 cell lines compared to vehicle-treated control cells than either Omipalisib or Trametinib treatment alone (Figure 2c). OmiTram treatment in the K8484 and PKT62 cell lines suppressed colony-forming ability slightly more effectively, but did not reach significance (Figure 2c). Treatment with OmiSHP more effectively suppressed colony formation compared to vehicle-treated controls in the K8484 and MiaPaCa2 cell lines compared to either Omipalisib or SHP099 treatment alone. OmiSHP treatment trended toward suppressing colony formation more effectively in PKT62 and Panc1 cell lines compared to individual treatments, but the difference did not reach significance (Figure 2d). Overall, colony formation was significantly suppressed using OmiSHP for all cell lines when compared to vehicle.

We next measured the effectiveness of each combination treatment strategy on PDAC cell migratory ability using a scratch wound-healing assay (Figure 3). Both OmiTram and OmiSHP treatment more effectively suppressed cell migration compared to vehicle-treated control cells than either Omipalisib or Trametinib treatment alone (Figure 3a, b). However, when assessing combination treatment compared to each individual treatment, OmiTram treatment was not significantly more effective at blocking migration compared to either individual Omipalisib or Trametinib treatment (Figure 3a), while OmiSHP treatment significantly inhibited migration compared to Omipalisib treatment alone in the PKT cell line and the SHP099 treatment alone in the K8484, MiaPaCa2, and Panc1 cell lines (Figure 3b). Overall, combined inhibition of MAPK and PI3K pathways via Omipalisib/Trametinib or Omipalisib/SHP099 treatment *in vitro* suppresses PDAC tumor cell proliferation, migration, and colony forming ability.

**Figure 3.**
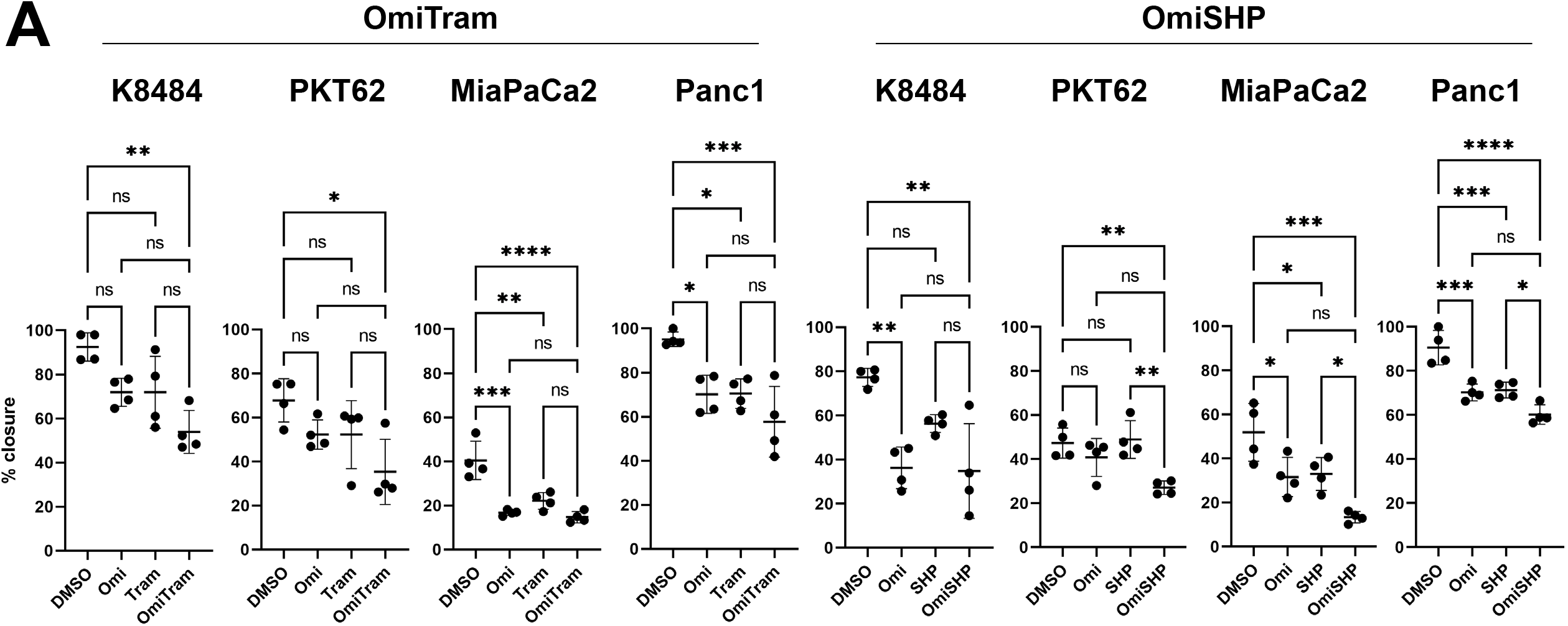
Combined targeting of PI3K and MAPK pathways inhibits migratory ability of PDAC cells *in vitro*. Changes in migratory ability after treatment with either OmiTram **(A)** or OmiSHP **(B)** was assessed via scratch migration assay. K8484 cells were treated with 5nM Omipalisib, 10nM Trametinib, and/or 50uM of SHP099 over 24 hours (OmiTram) or 48 hours (OmiSHP). PKT62 cells were treated with 10nM Omipalisib, 10nM Trametinib, and/or 50uM of SHP099 over 8 hours. MiaPaCa2 cells were treated with 25nM Omipalisib, 20nM Trametinib, and/or 20uM SHP099 over 48 hours. Panc1 cells were treated with 50nM Omipalisib, 20nM Trametinib, and/or 50uM SHP099 over 24 hours. n=4 wells for all groups. (*p < 0.05, **p <0.01, ***p < 0.001, and ****p < 0.0001).

### Combined MAPK and PI3K pathway inhibition suppresses tumor growth *in vivo*

We next assessed the efficacy of a combined MAPK-PI3K targeted therapeutic strategy *in vivo*. First, to monitor the effect of combination MAPK and PI3K inhibition on tumor growth *in vivo* over time, we injected PDAC tumor cells (K8484) subcutaneously into the flank of C57BL/6 mice. After tumors reached a sufficient starting volume, mice were treated 3 times per week via oral gavage with either vehicle control (methylcellulose), Omipalisib (0.3 mg/kg), Trametinib (1 mg/kg), SHP099 (50mg/kg), combined Omipalisib/Trametinib (OmiTram), or combined Omipalisib/SHP099 (OmiSHP). Tumor volume was measured every 2 days for 18 days. Overall, all therapeutic treatments suppressed tumor growth compared to the vehicle control. Over the 18 day study period, the tumor growth rate of the Trametinib (alone), OmiTram, and OmiSHP groups were significantly suppressed compared to the control group (Figure 4a). However, while the OmiSHP or OmiTram-treated mice displayed no net overall tumor growth increase for the first 10 days of the study, the OmiSHP group displayed tumor growth after 10 days resulting in more overall tumor growth compared to the OmiTram group by the 18-day study period endpoint (p=0.06) (Figure 4a). At day 18, the OmiTram treated mice represented the only group with both significantly reduced tumor size and weight compared to the vehicle control group (Figure 4a, b). Together, these data indicate that combined MAPK and PI3K inhibition via OmiSHP or OmiTram represent therapeutic strategies to slow tumor growth *in vivo*, but an Omipalisib/Trametinib combined therapeutic effectively inhibited tumor growth over time.

**Figure 4.**
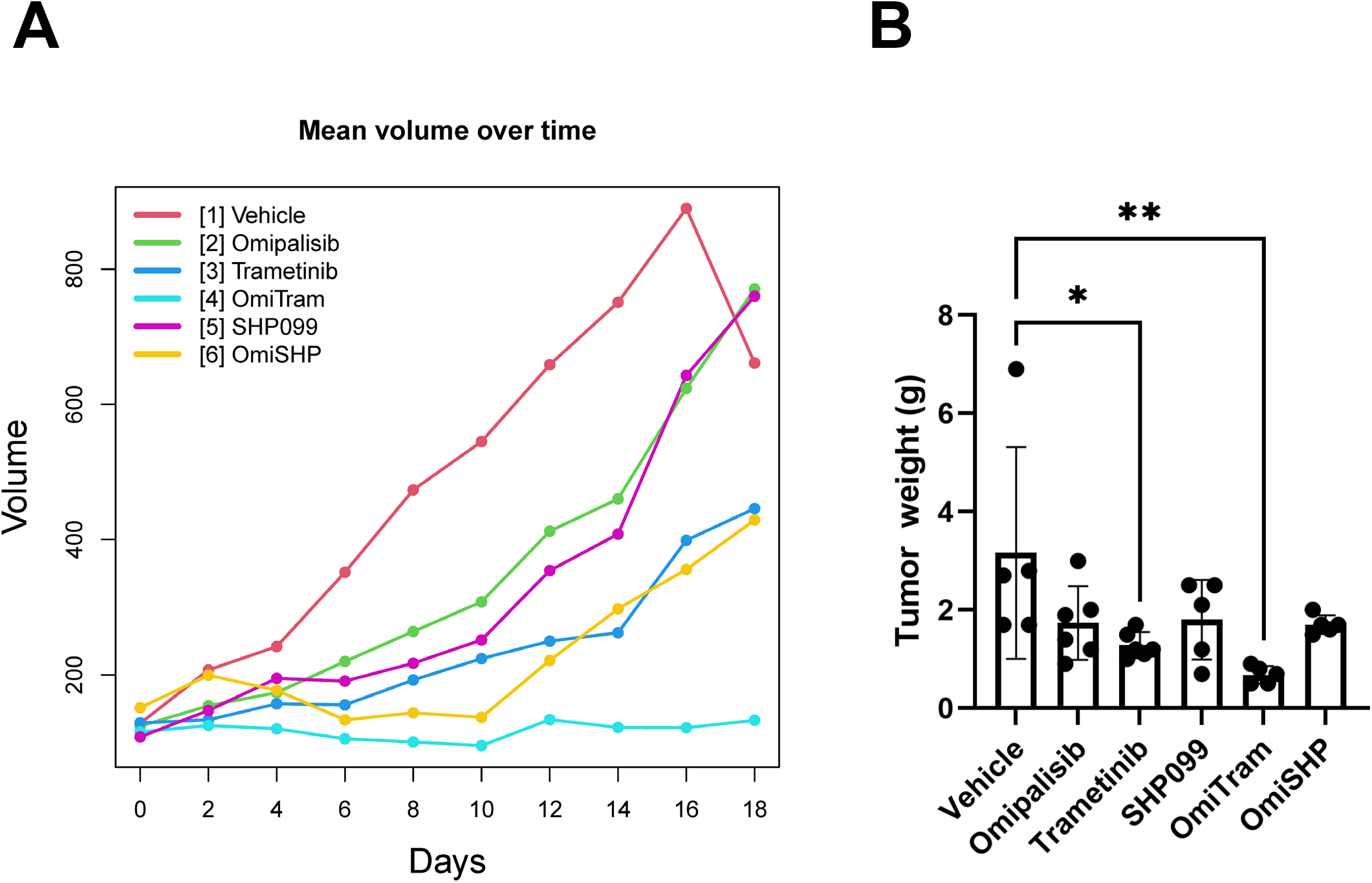
Combined targeting of PI3K and MAPK pathways suppresses tumor growth in an *in vivo* subcutaneous model of PDAC. **(A)** Mean tumor volume over time of mice treated with Omipalisib, Trametinib, SHP099, combination OmiSHP or combination OmiTram targeted therapeutic. **(B)** Tumor weight analysis after 18 days of treatment. Vehicle: n=5; Omipalisib: n=6, Trametinib: n=6, SHP099, n=6; OmiTram, n=5; OmiSHP, n=6. (*p < 0.05, **p <0.01, ***p < 0.001, and ****p < 0.0001).

The OmiTram combination therapeutic treatment resulted in the most effective inhibition of tumor growth in a subcutaneous model of PDAC, therefore we further assessed whether the OmiTram administration would also suppress tumor growth and progression in an aggressive spontaneous mouse model of PDAC. The PKT (PTf1a^cre/+^;LSL-^KRASG12D/+^;TGFBR2^flox/flox^) mouse model typically exhibits PanIN formation at 3.5 weeks of age and reaches endpoint at 7-10 weeks of age (49). PKT mice received vehicle control (methylcellulose), Omipalisib (0.3 mg/kg), Trametinib (1 mg/kg), or OmiTram treatment 3 times per week starting at 4.5 weeks of age and continuing until humane endpoint was reached or mice surpassed 14 weeks of age (10 weeks of treatment was completed). While all of the mice in the Omipalisib and Trametinib single treatment groups reached humane endpoint before the experimental endpoint, 5 of 7 mice treated with the OmiTram combination therapeutic completed 10 weeks of treatment and surpassed 14 weeks of age, more than doubling the average survival time of the vehicle-treated mice (figure 5a). Tumor weight normalized to total body weight at endpoint was significantly lower in the OmiTram group than the vehicle, Omipalisib, and Trametinib groups (Figure 5b). Additionally, pancreata from the OmiTram group exhibited significantly less tumor area than the Omipalisib or Trametinib groups compared to vehicle (p=0.008) (Figure 5c). Overall, these data indicate that a simultaneous targeting of PI3K and MAPK signaling pathways via Omipalisib and Trametinib more effectively suppresses tumor growth than treatment with either individual drug.

**Figure 5.**
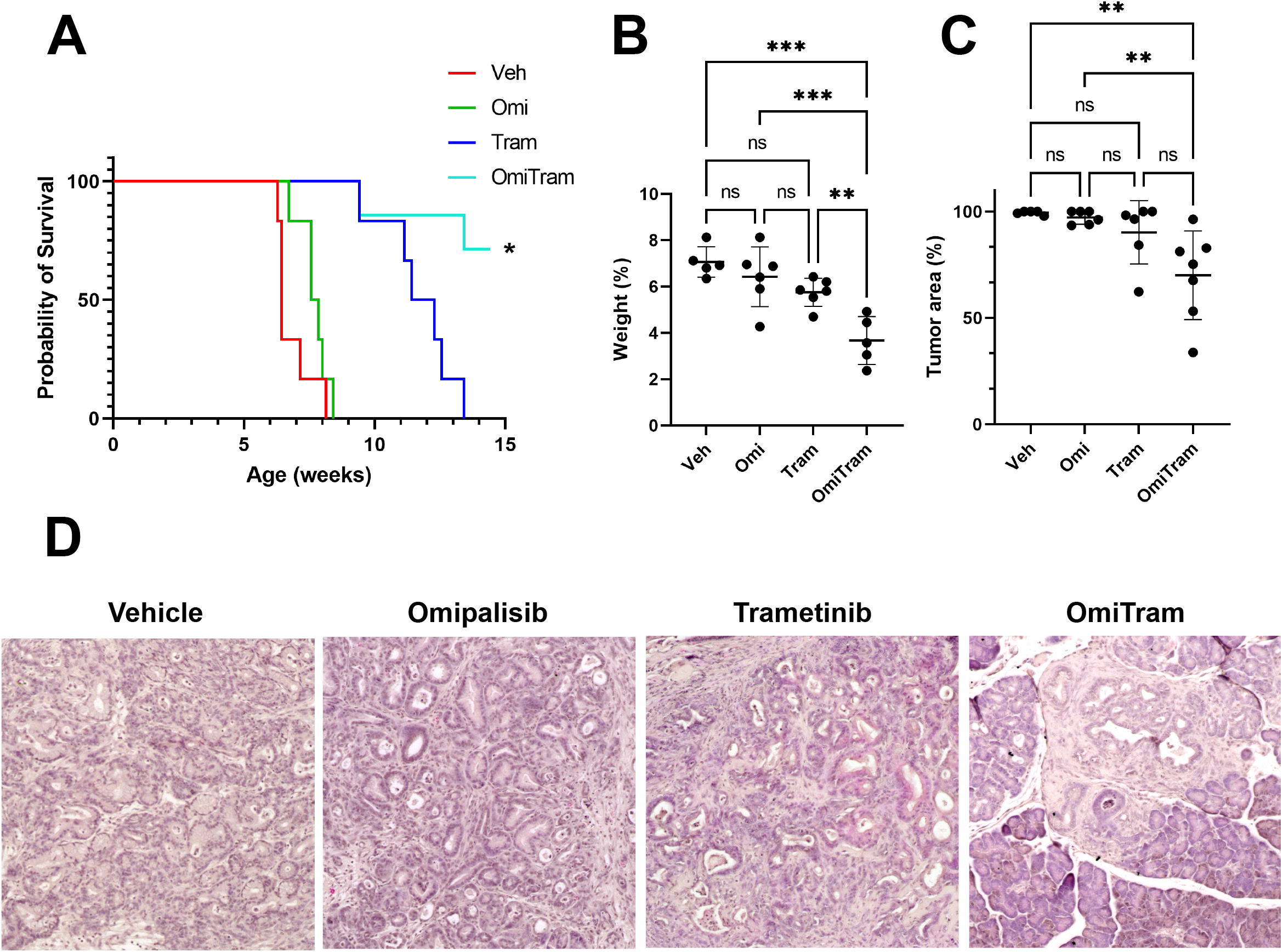
Combined targeting of PI3K and MAPK pathways suppresses tumor growth in an *in vivo* spontaneous mouse model of PDAC. **(A)** Survival analysis of mice treated with vehicle, Omipalisib, Trametinib, or combined OmiTram targeted therapeutic. * represents the study endpoint (10 weeks after start of treatment, or 14.5 weeks of age). Vehicle: n=6; Omipalisib: n=6; Trametinib: n=6; OmiTram: n=7. **(B)** Endpoint tumor weight was measured as a percentage of total mouse weight. **(C)**. % tumor area was measured from H&E-stained tissue sections (representative H&E images in **(D)**). Post mortem analysis: vehicle: n=5; Omipalisib: n=6; Trametinib: n=5; OmiTram n=6. (*p < 0.05, **p <0.01, ***p < 0.001, and ****p < 0.0001).

## Discussion

Our study has demonstrated sustained or upregulated PI3K pathway activation upon MAPK pathway inhibition with Trametinib or SHP099 as well as sustained or upregulated MAPK pathway activation upon PI3K/mTOR inhibition via Omipalisib in PDAC. These results support others that show upregulation of MAPK activation upon inhibition of various components of the PI3K-AKT-mTOR pathway in solid cancers and vice versa (26, 41, 50). Inversely, others have shown that PI3K inhibition alone also induces MAPK pathway inhibition in cancer rather than resulting in compensatory activation (19, 51, 52). Simultaneous pERK inhibition upon PI3K inhibition has also been shown in PDAC tumor cells, but in the presence of wild-type, rather than mutant, KRAS (19). Since the vast majority of PDAC presents with mutant KRAS (5-8), dual therapeutics targeting PI3K and direct inhibitors of mutant KRAS (13, 53) may prove more synergistic than therapeutics targeting downstream targets of mutant RAS (19, 26).

Awasthi et al. and Junttila et al. et al have shown that, in preclinical models, combined MAPK and PI3K inhibition enhance PDAC tumor response to Gemcitabine and Gemcitabine/nab-paclitaxel, the current standard chemotherapeutic strategies (54, 55). However, the effect was modest and, in the case of combined MAPK/PI3K inhibition combined with gemcitabine alone, this treatment strategy did not show more effectiveness than combined gemcitabine/erlotinib treatment, which is already approved in the clinic (55). Toxicity was identified as a major consideration in combination therapeutic strategies involving MAPK and PI3K inhibition. Though dual inhibition of MEK and PI3K has shown promise in pre-clinical models, it has not been well-tolerated in clinical trials involving patients with advanced solid tumors, including PDAC (56, 57). For example, a phase 1b clinical trial using a combined Omipalisib/Trametinib strategy demonstrated some antitumor efficacy but was ultimately terminated due toxicity (57). Though SHP2 inhibitors are being investigated in clinical trials (NCT03114319, NCT05354843) and SHP2 inhibition has been preliminarily shown to counteract therapeutic resistance to MEK inhibition (15), further investigation of whether a SHP2/PI3K-targeted dual therapeutic strategy would demonstrate clinical efficacy might be considered.

Altogether, our results demonstrate that MAPK inhibition via Trametinib or SHP099 combined with PI3K/mTOR inhibition via Omipalisib act synergistically to inhibit PDAC tumor cell growth and migration more effectively than either single therapeutic agent alone. While an Omipalisib/Trametinib therapeutic strategy was more effective than an Omipalisib/SHP099 therapeutic, a clinical trial has already demonstrated significant toxicity of an Omipalisib/Trametinib using an effective doses. Therefore, further technological developments are needed to develop tumor selectively or targeted delivery to achieve optimal combination therapeutic strategies to synergistically target MAPK and PI3K pathways in PDAC.

## Acknowledgements

Research reported in this publication was supported by the National Cancer Institute Cancer Center Support Grant P30 CA168524 (PI: Jensen) CCSG pilot funding for Dr. VanSaun. We acknowledge the Flow Cytometry Core Laboratory, which is sponsored, in part, by the NIH/NIGMS COBRE grant P30 GM103326 and the NIH/NCI Cancer Center grant P30 CA168524. The K8484 cell line was provided by Dave Tuveson lab characterized in Olive et al. 2009 (46). The PKT62 cell line was provided by Nagaraj Nagathihalli (University of Miami).

## Author Contributions

Conception and design: B.A. Bye, J. Jack, M.N. VanSaun

Development of methodology: B.A. Bye, J. Jack, M.N. VanSaun

Acquisition of data: B.A. Bye, J. Jack, A. Pierce, R.M. Walsh, A. Olou

Analysis and interpretation of data (e.g., statistical analysis, biostatistics, computational analysis): B.A. Bye, J. Jack, A. Pierce, R.M. Walsh, A. Olou, P. Chalise, M.N. VanSaun

Writing, review, and/or revision of the manuscript: B.A. Bye, J. Jack, A. Pierce, R.M. Walsh, P. Chalise, A. Eades, A. Olou, M.N. VanSaun

Administrative, technical, or material support (i.e., reporting or organizing data): B.A. Bye, J. Jack, A. Pierce, R.M. Walsh, P. Chalise, A. Olou, M.N. VanSaun

Study supervision: M.N. VanSaun

## Notes

**Competing Interests Statement:** This work was supported in part by grant R01 CA231052 from the NCI to MNV, as well as by funds from the University of Kansas Cancer Center.

### Competing Interest Statement

The authors have declared no competing interest.

