## Supplemental files for "Combined PI3K and MAPK inhibition synergizes to suppress PDAC"

**Supplemental figure 1. Omipalisib dose-response assay in human and mouse PDAC cell lines.** PI3K pathway inhibition (as measured by pAKT levels) and consequent changes in MAPK pathway activity (as measured by pERK) was assessed in 2 mouse (K8484 and PKT62) and 2 human (MiaPaCa2 and Panc1) PDAC cell lines. Cells were treated for 24 hours with Omipalisib before lysis.

Supplemental Figure 1

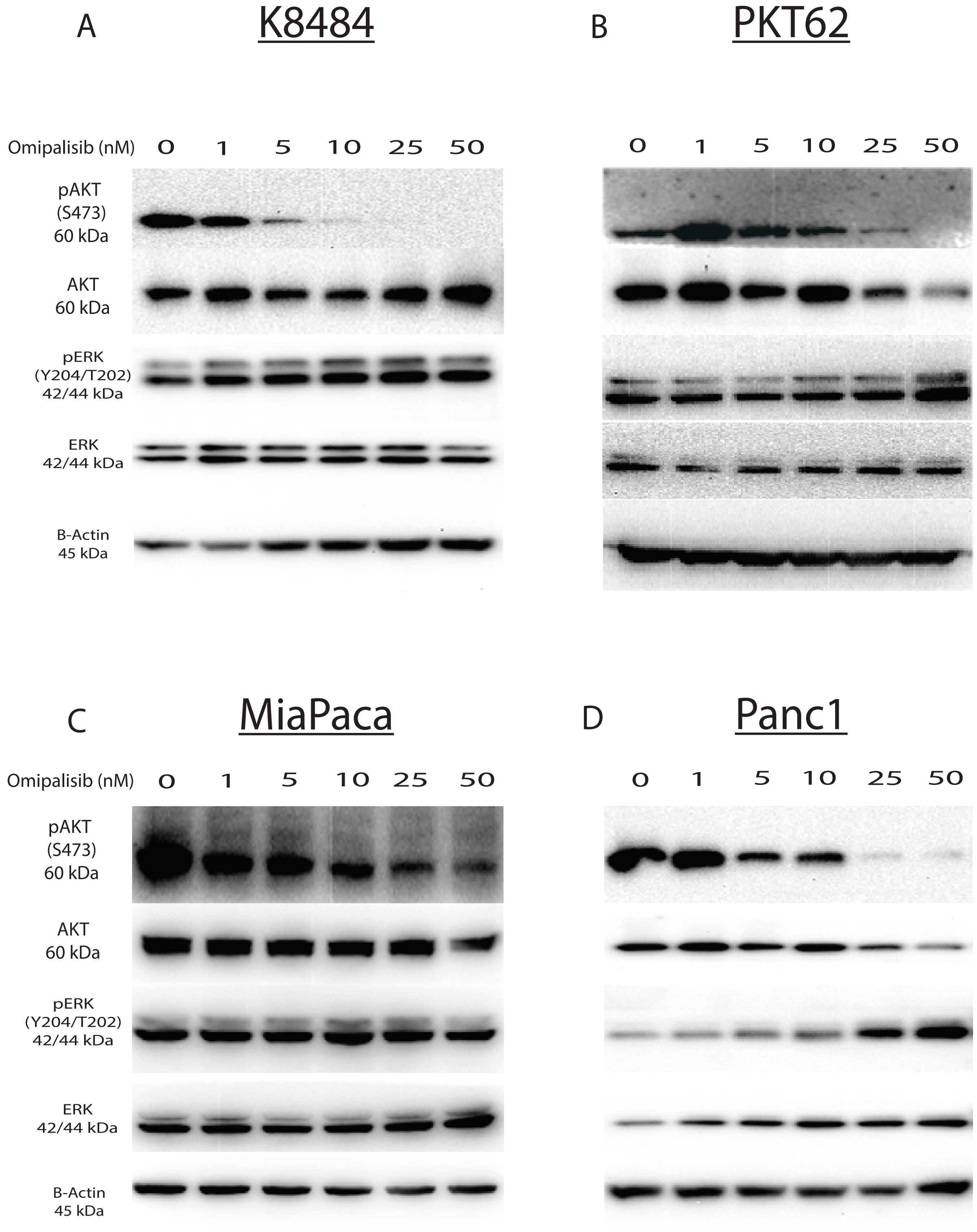

**Supplemental figure 2. Trametinib dose-response assay in human and mouse PDAC cell lines.** MAPK pathway inhibition (as measured by pERK levels) and consequent changes in PI3K pathway activity (as measured by pAKT) was assessed in 2 mouse (K8484 and PKT62) and 2 human (MiaPaCa2 and Panc1) PDAC cell lines. Cells were treated for 24 hours with Trametinib before lysis.

Supplemental Figure 2

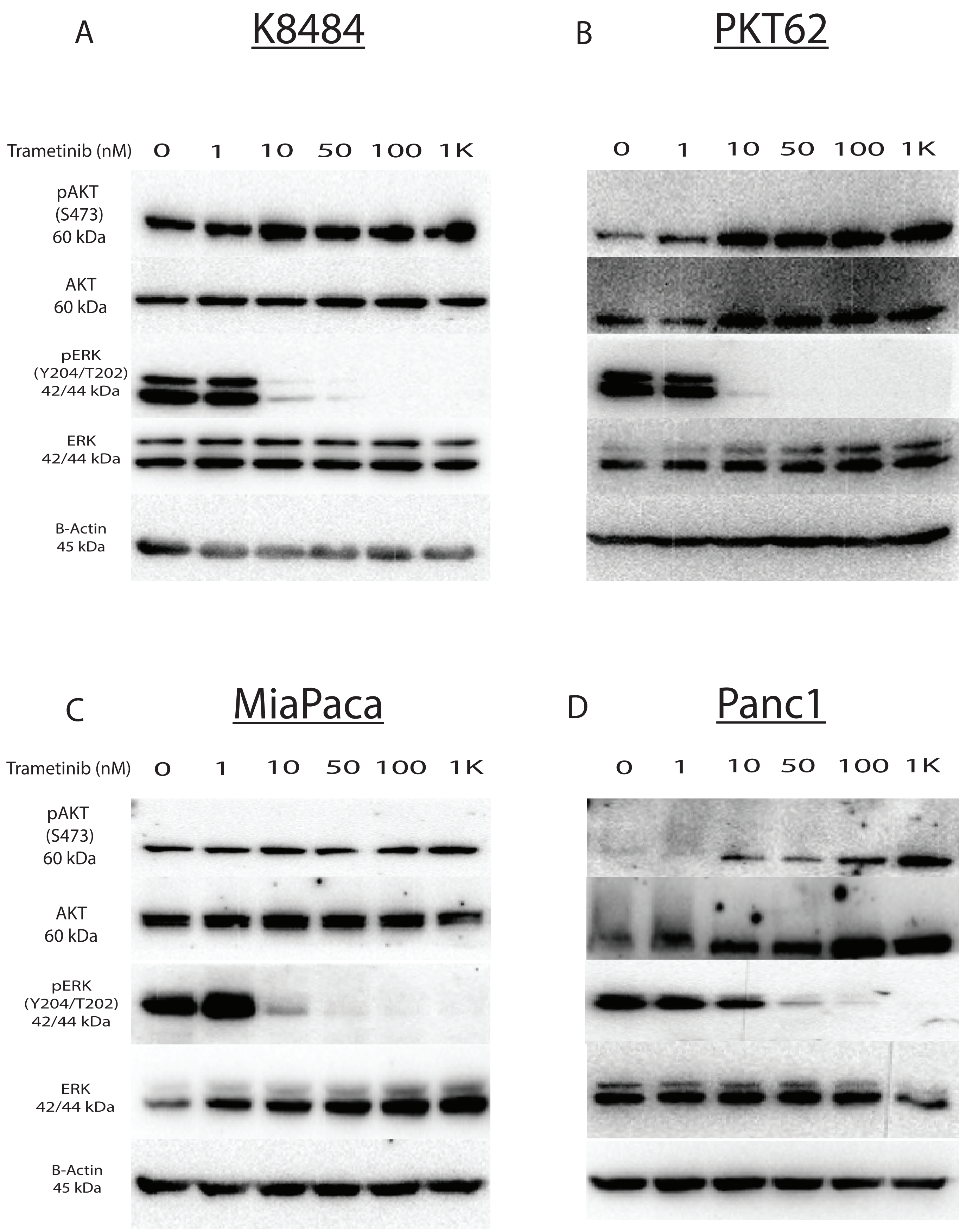

**Supplemental figure 3. SHP099 dose-response assay in human and mouse PDAC cell lines.** MAPK pathway inhibition (as measured by pERK levels) and consequent changes in PI3K pathway activity (as measured by pAKT) was assessed in 2 mouse (K8484 and PKT62) and 2 human (MiaPaCa2 and Panc1) PDAC cell lines. Cells were treated for 3 hours with SHP099 before lysis.

Supplemental Figure 3

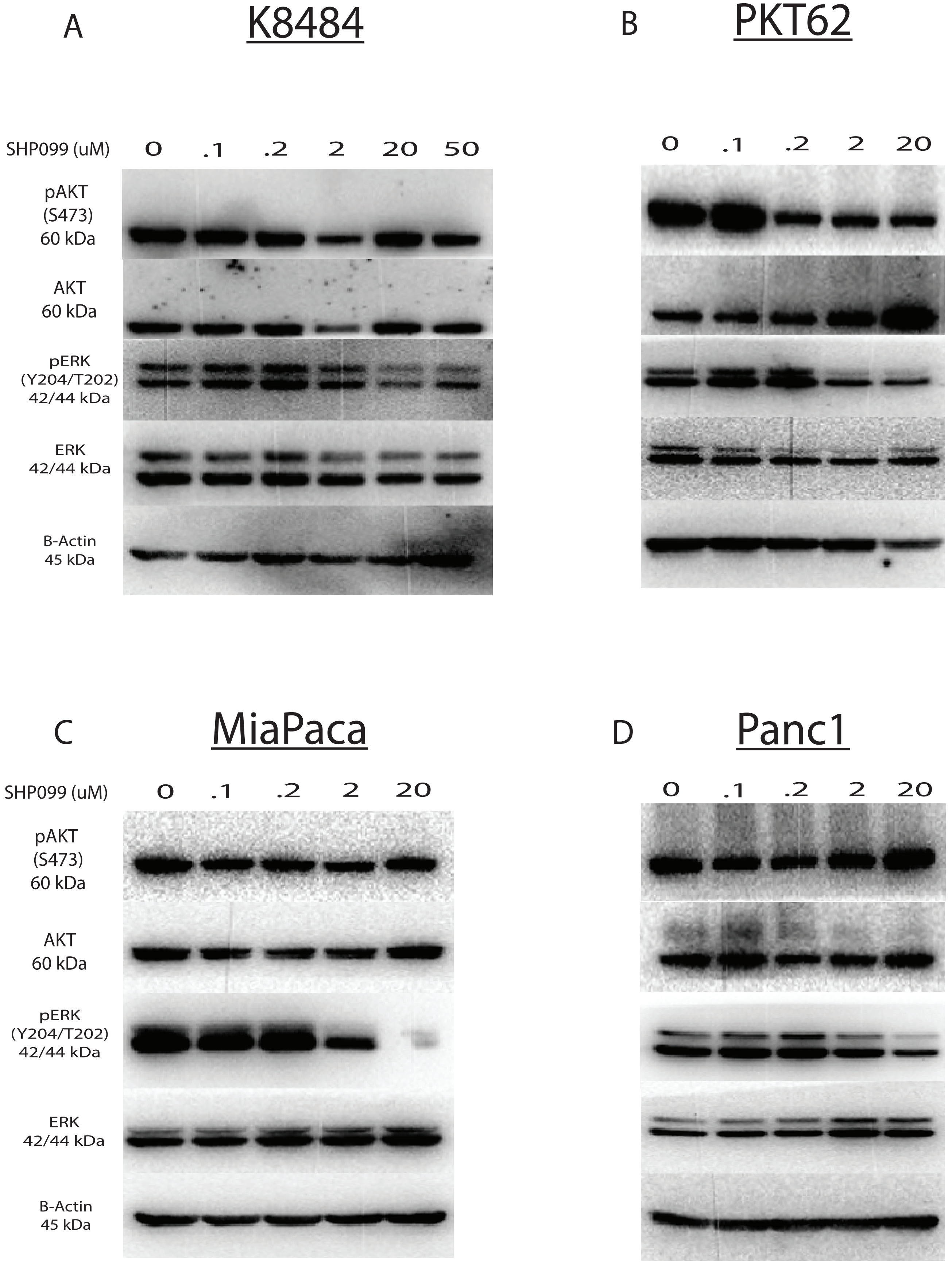
